# A scoping review on the use of consumer-grade EEG devices for research

**DOI:** 10.1101/2022.12.04.519056

**Authors:** Joshua Sabio, Nikolas S Williams, Genevieve M McArthur, Nicholas A Badcock

## Abstract

**BACKGROUND:** Commercial electroencephalography (EEG) devices have become increasingly available over the last decade. These devices have been used in a wide variety of fields ranging from engineering to cognitive neuroscience.

**PURPOSE:** The aim of this study was to chart peer-review articles that used currently available consumer-grade EEG devices to collect neural data. We provide an overview of the research conducted with these relatively more affordable and user-friendly devices. We also inform future research by exploring the current and potential scope of consumer-grade EEG.

**METHODS:** We followed a five-stage methodological framework for a scoping review that included a systematic search using the Preferred Reporting Items for Systematic Reviews and Meta-Analyses Extension for Scoping Reviews (PRISMA-ScR) guidelines. We searched the following electronic databases: PsycINFO, MEDLINE, Embase, Web of Science, and IEEE Xplore. We charted study data according to application (BCI, experimental research, validation, signal processing, and clinical) and location of use as indexed by the first author’s country.

**RESULTS:** We identified 916 studies that used data recorded with consumer-grade EEG: 531 were reported in journal articles and 385 in conference papers. Emotiv devices were most used, followed by the NeuroSky MindWave, OpenBCI, interaXon Muse, and MyndPlay Mindband. The most common use was for brain-computer interfaces, followed by experimental research, signal processing, validation, and clinical purposes.

**CONCLUSIONS:** Consumer-grade EEG has proven to be a useful tool for neuroscientific research and will likely continue to be used well into the future. Our study provides a comprehensive review of their application, as well as future directions for researchers who wish to use these devices.

## Introduction

Electroencephalography (EEG) is the continuous measurement of electrical activity generated by neurons firing in the brain. This involves placing metal electrodes at multiple sites on the scalp record fluctuations in voltage at millisecond-level temporal resolution. Such recordings can then be processed to produce spectral analyses of electrical activity or generate event-related potentials (ERPs) that represent the averaged response to a stimulus. Today, EEG is one of the most popular neuroscientific tools for academics and medical professionals due to its non-invasiveness and ease-of-use [1]. In recent years, several companies have developed consumer-grade EEG devices. These devices are compact, wireless, and have a streamlined setup, making them particularly attractive to novice researchers or those looking to collect data outside the traditional laboratory setting [2]. More importantly, consumer-grade devices are cheaper than research-grade devices, allowing those with limited funding an affordable means to collect EEG data.

Due to its accessibility, consumer-grade EEG has been used for a wide variety of purposes across different fields. Software engineers and computer scientists use consumer-grade EEG to collect high resolution time-series data. This data is then processed to create or optimise machine learning and signal processing algorithms [3–5]. In turn, these algorithms can be used in conjunction with the device to develop brain-computer interface (BCI) systems. Those in engineering and robotics can train machines to respond in real time to patterns found in neural data [6]. Once synchronised, a human user can configure a BCI to control a multitude of electronic devices including wheelchairs [7], drones [8], smart homes [9–11], and web browsers [12]. Clinicians report using the technology to administer neurofeedback therapy [13], facilitate learning [14], assess patient sleep quality [15,16], and determine affective states [17–20]. Scientists increasingly use consumer-grade devices to collect neural data to address a variety of theoretical and practical research questions [2,21,22].

The proliferation of research with consumer-grade EEG has inspired several non-systematic reviews (see Table 1). For instance, some reviews compared the performance of a single consumer-grade EEG device to non-EEG biosensors in the domains of seizure detection [23], BCI systems [24], and stress recognition [25]. Others have compared *multiple* consumer-grade EEG devices within a single domain [2,21,26–28]. One of the most thorough of these reviewed around 100 “handpicked” [p. 3997] studies that used four consumer-grade devices – the NeuroSky MindWave, Emotiv EPOC+, interaXon Muse, and OpenBCI neuroheadset – in the domains of “cognition, brain-computer interfaces (BCI) [sic], education research, and game development” [22].

**Table 1.** List of literature reviews that include 92 consumer-grade EEG devices.

| First Author | Year | Domain | Devices Reviewed |
| --- | --- | --- | --- |
| Rechy-Ramirez et al. | 2017 | BCI | Emotiv |
| Xu et al. | 2018 | Educational research | Emotiv; NeuroSky MindWave |
| Aldridge et al. | 2019 | P300 speller task | OpenBCI |
| Kurada et al. | 2019 | Seizure detection | Emotiv |
| Roy et al. | 2019 | Deep learning | Emotiv; InteraXon Muse; Myndplay Mindband; NeuroSky MindWave; OpenBCI |
| LaRocco et al. | 2020 | Drowsiness detection | Emotiv; InteraXon Muse; NeuroSky MindWave; OpenBCI |
| Vasiljevic et al. | 2020 | BCI gamification | Emotiv; Myndplay Mindband; NeuroSky MindWave; OpenBCI |
| Shum et al. | 2020 | Seizure detection | Emotiv; NeuroSky MindWave |
| Castro-Garcia et al. | 2021 | Stress recognition | OpenBCI |

*Note.* These reviews were excluded from the present review for reasons outlined in Method and Results.

While these non-systematic reviews provide insights into the domain-*specific* functions of certain EEG devices, the current literature on this topic is, at best, fragmented. Indeed, it is surprising that, to date, there has been no systematic review on the research-related use of currently available and commonly used consumer-grade EEG devices. Thus, the aim of this paper is to synthesise the large volume of studies that have used consumer-grade EEG to collect neural data. We categorise each of these studies under an appropriate domain or category to provide a domain-*general* snapshot of the exciting work conducted with this emerging technology.

## Method and Results

We followed the Preferred Reporting Items for Systematic Reviews and Meta-analyses Extension for Scoping Reviews (PRISMA-ScR) guidelines [29]. Scoping reviews report the “extent, range, and nature of research activity” [p. 21] in five stages: (1) identify the research question; (2) identify relevant studies; (3) select the studies; (4) chart the data; (5) collate, summarise, and report the results.

### Step 1: Identify the research question

We aimed to explore the extent to which consumer-grade EEG have been applied to research domains in locations around the world. Regarding EEG devices, our goal was to identify the EEG systems produced by companies that manufacture the most popular consumer-grade devices. Regarding research domains, our goal was to identify the primary research fields that these consumer-grade EEG systems have been applied to. Finally, regarding research location, our goal was to answer the question of where the devices have been used globally.

### Stage 2: Identifying relevant studies

We conducted a systematic literature search by retrieving records from five online bibliographic databases: (a) PsycINFO, (b) MEDLINE, (c) Embase, (d) Web of Science, and (e) IEEE Xplore. These databases contain studies from multiple fields including neuroscience, psychology, medicine, and engineering. Searches included studies published from 2010, as this is the year the first studies examining consumer-grade EEG were published. Studies also had to be conducted with human participants and written in English, which is currently the universal scientific language. To find records in each database, we used search strings to capture keyword variations (see Table 3). These were subsequently adjusted to fit the search syntax of each database.

### Stage 3: Study selection

The initial complete search was conducted in April 2022 and yielded 1260 articles. We excluded 265 duplicates and screened the remaining. After screening, we excluded a further 79 studies as they met one or more of the following exclusion criteria: no device was used, or no EEG was data collected (n = 53); not written in English (n = 6); had inaccessible PDFs (n = 20). The final number included was 916, comprising 531 journal articles and 385 conference proceedings. Figure 1 depicts the flow chart of the screening process.

**Fig 1.**
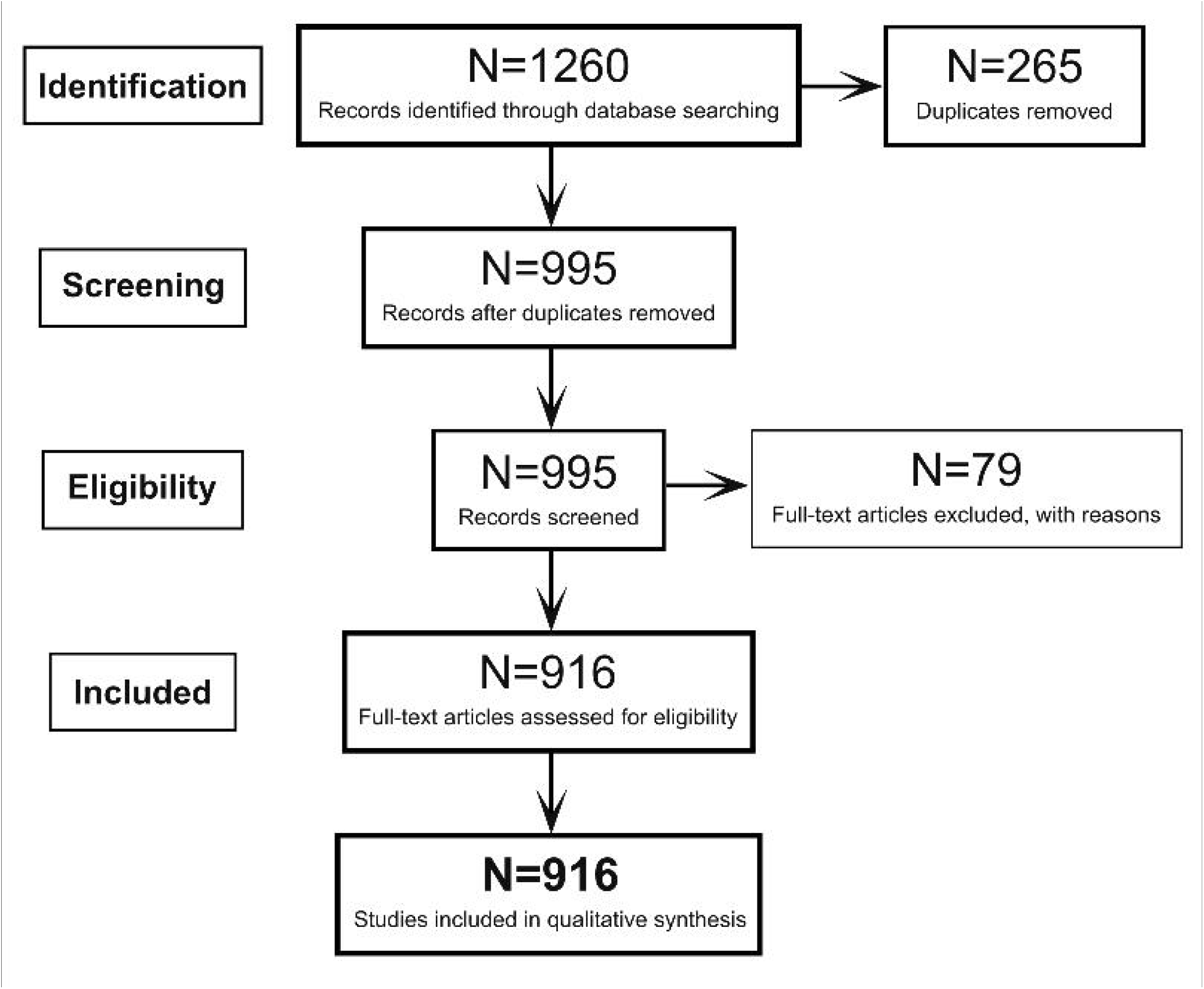
PRISMA flowchart of the screening process.

### Stage 4: Charting the data

We charted 916 included articles by recording the following relevant information: EEG device, research domain, authors, first author’s country, year of publication. Country information was obtained from the first author’s affiliation as listed in the article. If this could not be found, the first author’s most recent affilation was used.

### Stage 5: Collating, Summarising, and Reporting the Results Consumer-grade EEG devices

The devices used by articles included in this scoping review were produced by five companies: **Emotiv** (67.69%), **NeuroSky MindWave** (24.56%), and **OpenBCI, interaXon**, and **MyndPlay** (7.75% collectively for these three devices). **Emotiv** has released three versions of a 14-channel EEG device: EPOC, EPOC+, and EPOC-X. For simplicity, all three devices will hereon be referred to as “Emotiv.” The EPOC+ and EPOC-X can capture data at both 128 and 256 Hz sampling rates, whereas EPOC captures data at 128 Hz. The EPOC-X includes an updated amplifier configuration, sensor housing, and rotating headband. **OpenBCI** provides two highly customisable EEG systems: the four-electrode Ganglion board and the eight-electrode Cyton board. Again, for simplicity, both boards will be referred to as “OpenBCI.” Users can configure an assembly of eight electrodes around an OpenBCI Cyton processing circuit board and daisy-chain it to another Cyton board for a 16-channel system. The **interaXon Muse** is a headband-shaped device that records at 256 Hz from four electrodes located above the eyes (AF7, AF8) and above the ears (TP9, TP10). The **MyndPlay MyndBand** records at 512 Hz from three electrodes on the forehead. The **NeuroSky MindWave** is shaped like a gaming headset, records at 512 Hz from a single-channel electrode on the forehead (FP1), and pairs with a mobile application that can detect power spectra and other metrics such as meditation and attention using built-in signal processing techniques.

Research and consumer-grade EEG typically differ in their electrode count. Whereas research-grade EEG typically comprise dozens of electrodes fitted over the scalp with saline gel or solution, consumer-grade EEG have fewer electrodes and use minimal or no conductive medium. However, despite their low electrode count, consumer-grade EEG devices have been validated for scientific research in a broad range of paradigms, as outlined under Validation below. Table 2 lists the technical specifications of all currently available devices.

**Table 2.** Technical specifications of all currently available commercial EEG devices.

| Manufacturer | Model | Release Year | Channels | Sample rate (Hz) | Resolution |
| --- | --- | --- | --- | --- | --- |
| Emotiv | EPOC+ | 2013 | 14 | 128 or 256 | 14-bit |
| MyndPlay | Mindband | 2014 | 1 | 512 | * |
| OpenBCI | Ganglion board | 2014 | 4 | 200 | 24-bit |
| OpenBCI | Cyton board | 2014 | 8 | 250 | 24-bit |
| OpenBCI | Cyton-Daisy board | 2014 | 16 | 125 | 24-bit |
| Emotiv | Insight | 2015 | 5 | 128 | 14-bit |
| InteraXon | Muse 2 | 2016 | 4 | 256 | 12-bit |
| NeuroSky | MindWave Mobile 2 | 2018 | 1 | 512 | 12-bit |
| Emotiv | EPOC-X | 2019 | 14 | 128 or 256 | 14-bit |

*Note.* Emotiv also manufactures the EPOC Flex that contains 32 channels. This device is not included as it is explicitly marketed as a research device - not a commercial or consumer-grade device.

**Table 3.**
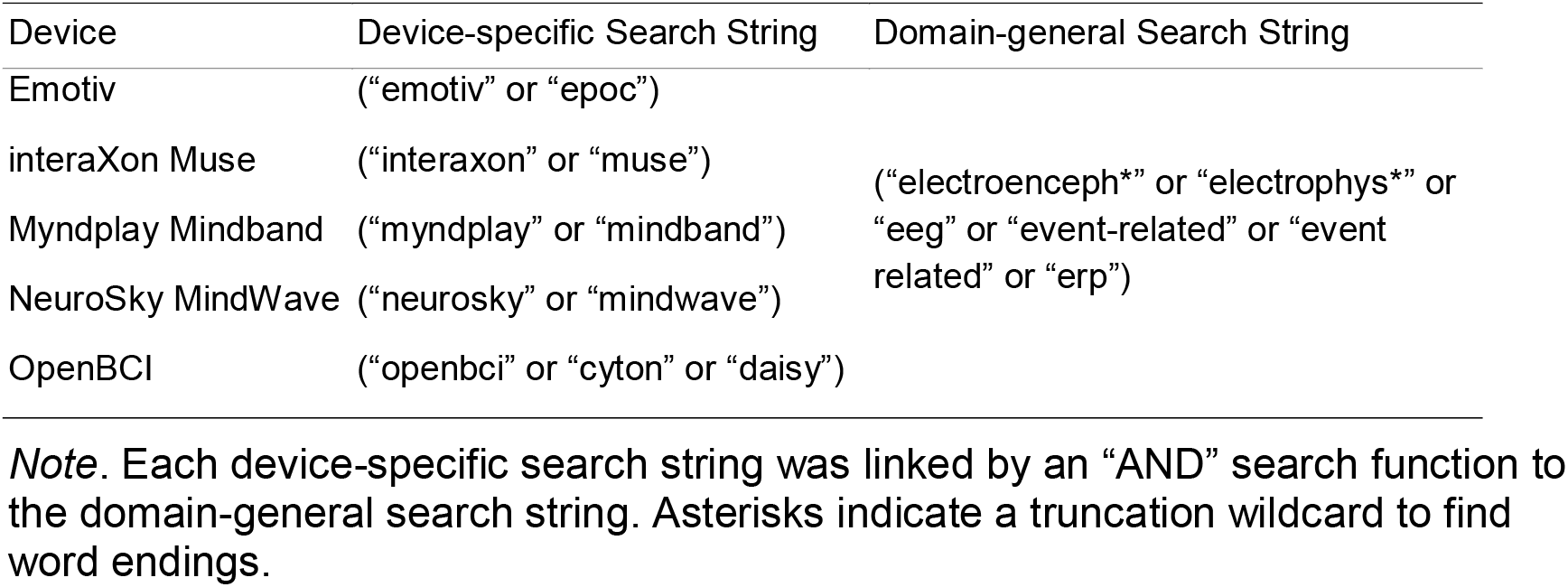
Search strings used for each device.

### Research domains

The included articles fell into five categories of use: BCI, experimental research, validation, signal processing, and clinical. Table 1 lists a description and an example for each usage category, and Figure 3 provides a visual depiction of device usage. It should be noted that these categories are not mutually exclusive. For example, a study that compared the performance of inexpensive, wireless, and dry EEG systems in classifying common neural responses [29] would be “validation.” In cases where a device could be classified under multiple categories, the authors made a joint decision about the appropriate category.

**Table 1.** Usage category descriptions and examples.

| Usage Category | Description | Example |
| --- | --- | --- |
| Brain-computer Interface (BCI) | Studies that facilitated interaction between human user and computer (or machine). | Realtime control of a wheelchair. |
| Experimental Research | Studies that used the device as a theory development tool | Examining EEG signatures in cognitive tasks. |
| Validation | Studies that aimed to validate use of the device | Comparing consumer-grade to research-grade EEG systems. |
| Signal Processing | Studies that collected data to refine EEG signals. | Creating a classification algorithm to reduce artefacts. |
| Clinical | Studies that used the device as a clinical tool | Assessing drowsiness in drivers. |

**Fig 2.**
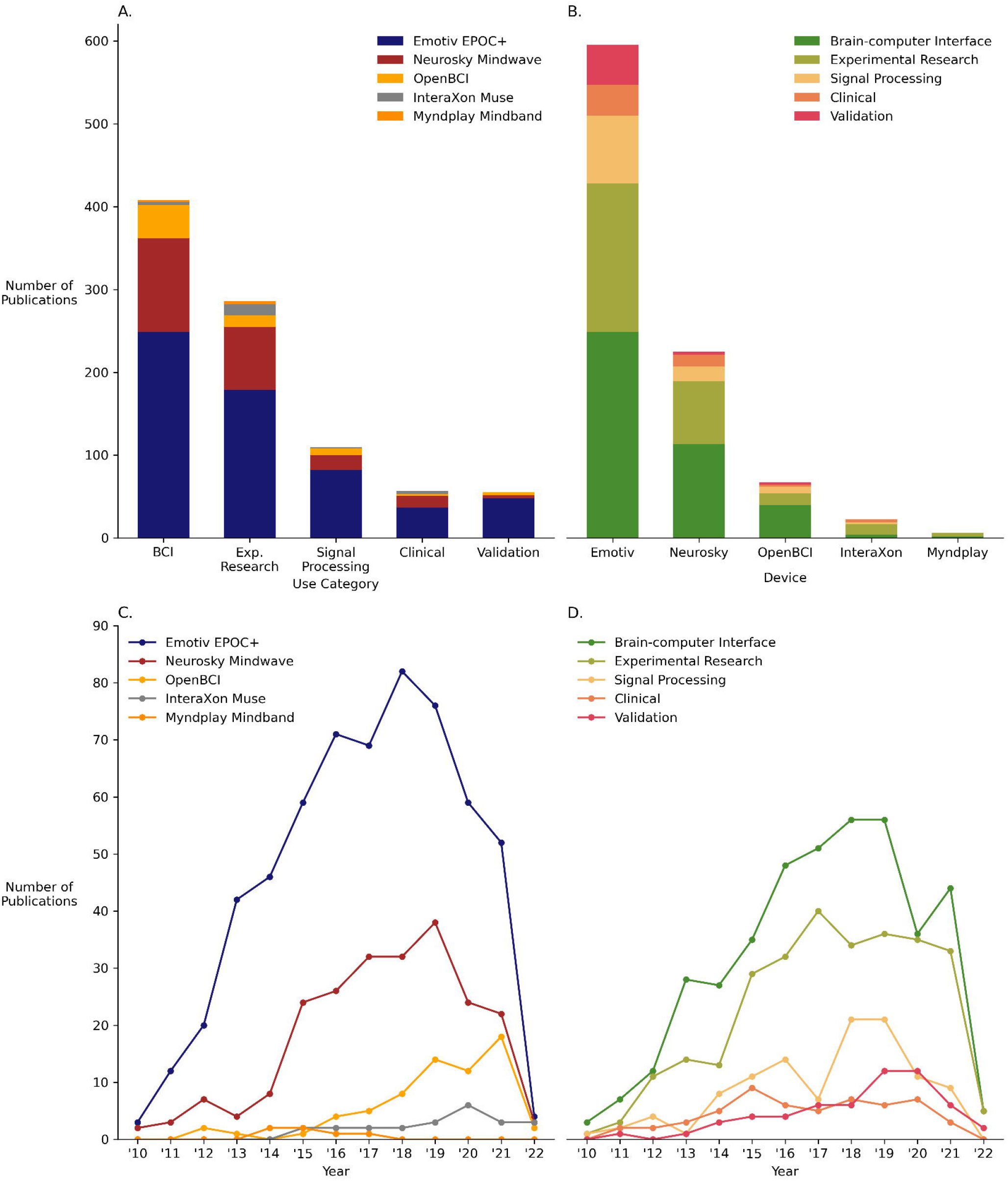
Charting the number of publications for each device and usage category. (A) Usage by device, collapsed. (B) Device by usage, collapsed. (C) Yearly publications for device (C). Yearly publications for usage (D)

**Fig 3.**
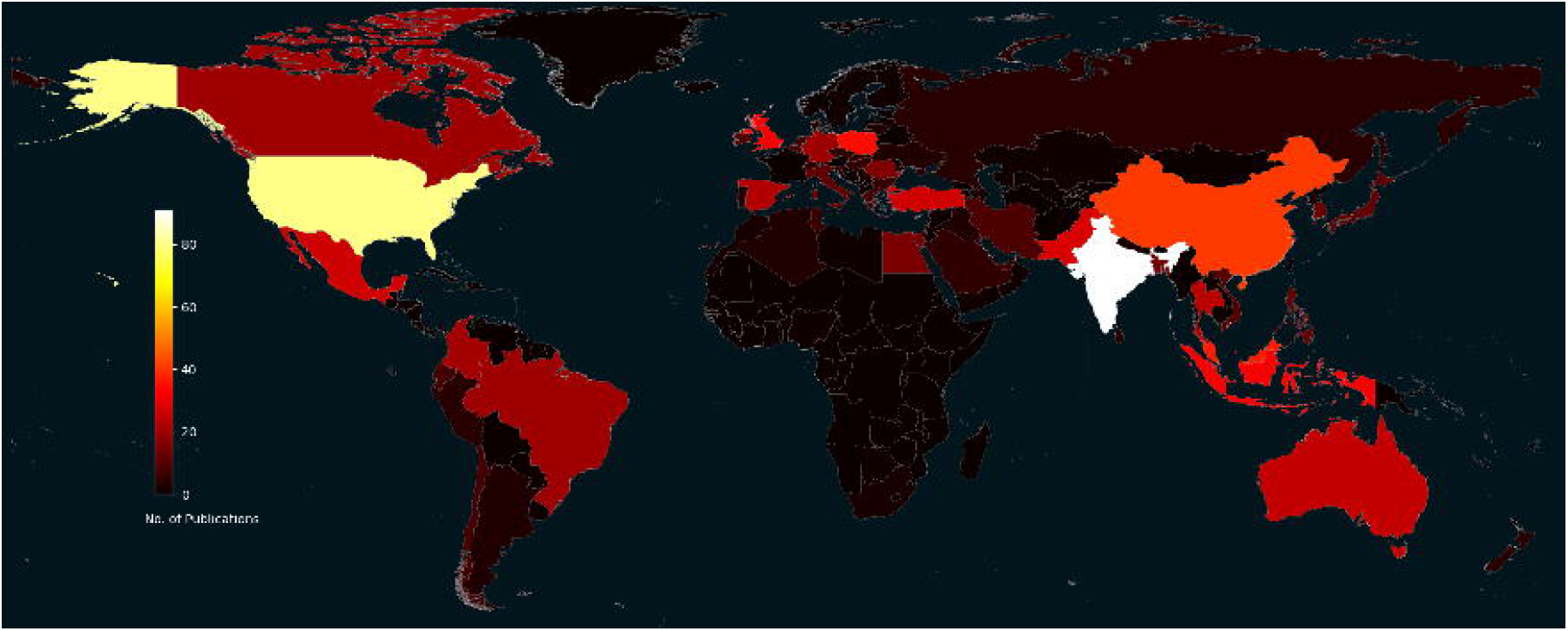
Geographical heatmap of publications that used consumer-grade EEG for research by country.

### Brain-computer Interfaces

The majority (44.54%) of included studies used consumer-grade EEG as an interface between a human user and computer. Through the application of machine learning algorithms, users are given the ability to control computers (software) or computer-based machines (hardware) and perform a variety of functions [27]. This is typically achieved in two steps. Firstly, data from a user is collected to train a classifier to recognise and extract features (i.e., spikes or longer pattern irregularities) from a continuous EEG signal, usually in response to some relevant stimuli. These features can then be evoked by the user to control a configured BCI system. How well the BCI system performs depends on multiple factors that include the chosen algorithm, the quality of the signal, electrode location, and the feature to be extracted [30–32].

The two earliest BCI-based studies were published in 2010. Using Emotiv, Ranky and Adamovich [33] trained participants over three weeks to move a robotic arm in three dimensions and grip certain objects using only facial movements. Likewise, Kwang-Ok et al. [34] used Emotiv to collect data for a paraplegic subject’s use of an emergency call system. These early BCI applications speak to the functional and clinical applicability of BCI systems, particularly for communication with and the motor improvement of paralysed patients.

One of the most common methods to configure a BCI system using consumer-grade EEG is with the steady-state visual evoked potential (SSVEP). SSVEPs are the typical neural response to visual stimuli that appear at matching frequencies [35]. In a recent study, Garcia et al. [36] used the interaXon Muse and SSVEPs (along with other signal processing algorithms) to decode and reconstruct stimuli in the visual field of participants with remarkable accuracy. In a similar study, Shi et al. [37] examined SSVEP image decoding applied to mobile phones. Chumerin et al. [38] designed a maze-navigating game that used SSVEPs and Emotiv; and together with an eye-tracker, Brennan et al. [39] tested the performance of an Emotiv in controlling a smart home with SSVEPs. Given the frequent use of SSVEPs, future researchers may also choose to take advantage of this feature’s high signal-to-noise ratio when implementing consumer-grade EEG into BCI systems.

Another easily identifiable ERP feature is the P300, a positive spike occurring roughly 300 ms after the presentation of an irregular, or “oddball”, stimulus in a stream of otherwise regular stimuli [40]. Due to its consistent latency, the P300 is commonly used as a target response in BCI-based applications [41]. Jijun et al. [42] used an Emotiv to capture P300 components and created a hands-free dialling system. Indeed, one of the most frequently observed applications of consumer-grade EEG is in P300 speller paradigms [43–47]. To illustrate, a matrix of letters is shown to an observer. To spell out a word, the observer must sequentially focus their attention on a single letter as each row and column in the matrix is rapidly highlighted. As one can imagine, P300 spellers prove extremely useful for those who have difficulties with verbal communication and expressive motor function – especially when combined with a portable device.

Motor-imagery is a purely mental process that involves the execution of motion without any explicit muscular or peripheral action [48]. There has been almost three decades of research involving BCI systems and motor imagery [49,50] – and their implementation with consumer-grade EEG is growing. By training an algorithm to detect specific EEG patterns reflecting imagined motor functions, users can operate artificial limbs [33,51–55] and wheelchairs [56–61]. Motor imagery with consumer-grade EEG has also been used to control electric vehicles [62] and aerial drones [8,63,64]. Parikh and George [65] designed a BCI system that uses motor imagery to control a quadcopter, and Das et al. [66] designed a BCI system to control a quadcopter with a user’s directional intentions.

Some of the most innovative BCI applications involved an interaction between a human user and software, rather than hardware. Shankar and Rai [67] used Emotiv to facilitate 3D computer-aided design (CAD) modelling, where user-evoked responses can activate various commands. Consumer-grade EEG has also been used to control web browsers [12]. Yehia et al. [68] designed an SSVEP-based website interface that presents options that can be selected with visual attention according to available functions on the current page.

More recent studies have explored the use of consumer-grade EEG in biometrics. Much like a fingerprint or password, algorithms can identify individuals based on neural patterns: Moctezuma et al. [69] identified participants via their EEG responses using imagined speech; Do et al. [70] identified participants according to their responses to images; and Saini et al. [71] collected responses while participants handwrote signatures to generate unique individual biomarkers.

### Experimental Research

Studies classified under “experimental research” used consumer-grade devices to collect EEG data outside of BCI applications. These studies comprised 31.22% of included articles and spanned a wide range of paradigms. For instance, the Emotiv was used in studies examining pilots’ reactions to unexpected events [72], reaction time [73], and mental fatigue and alertness [74]. Similarly, many experimental studies aimed to detect drowsy states [16,75–79]. Such paradigms have been adopted to improve road safety with the creation of early warning systems that alert a driver before an accident can occur [80–83]. These studies expand the application of EEG outside the laboratory for purely experimental purposes. Further studies have been conducted with consumer-grade EEG on students in the classroom [84–86]. Dikker et al. [87] used the Emotiv to measure the synchronised neural activity across students in a class, and found that this measure predicted overall classroom engagement and social dynamics.

Consumer-grade EEG has also been used to augment experiments testing athletes’ performance and mental states. Borisov et al. [88] used the Emotiv together with other biological indicators to examine athletes’ attention and stress levels. Liu et al. [89] correlated rifle-shooting accuracy with EEG signatures to find optimal states for good shots. Using the NeuroSky MindWave, Azunny et al. [90] found that meditation moderated athletes’ attention and working memory. Finally, Sultanov and Ismailova [91] determined power spectra during football players’ training and resting states.

While these studies report the performance of consumer-grade EEG across different experimental paradigms, we advise caution in applying consumer-grade devices to experiments that require excessive motion. EEG signals are inherently noisy. In cognitive and perceptual studies, researchers must remove trials where a participant’s movements introduce unwanted artefacts. In extreme cases, to avoid spurious results, entire participants are removed from analyses due to noisy artefacts. There are many in-house tutorials for popular toolboxes such as EEGLAB [92] and MNE [93], amongst other best-practice guidelines for cleaning EEG data [94–97]. We refer the reader to such guidelines to ensure the collection of high-quality neural signals for publication and replication.

### Signal Processing

Signal processing studies comprised 12.01% of included articles. These were studies that used consumer-grade EEG to create or improve signal processing algorithms or pipelines. Multiple studies have used an array of clustering and component analysis methods to remove eyeblinks and clean the signal recorded with consumer-grade EEG [98–105]. For instance, Szibbo et al. [106] proposed a novel algorithm to remove blink artefacts with logarithmic smoothing methods. Trigui et al. [107] used the Emotiv to accurately detect SSVEPs with the multivariate synchronisation index, an algorithm that characterises the synchronisation between recording and reference electrodes. Trigui et al. [108] also used the Emotiv to improve classification accuracy with a factor-analytic algorithm. Likewise, Elsawy et al. [109] proposed a principal components analysis classifier to use specifically on P300 spellers.

Given the relatively lower signal quality of consumer-grade relative to research-grade EEG, future research aiming to improve signal processing methods may apply these algorithms to data collected with research-grade EEG. This can assist further experimental EEG work by validating the algorithms’ robustness when applied to higher-quality data.

### Clinical

Clinical experiments comprised 6.22% of included articles. These studies involved the diagnosis and treatment of various physical and mental illnesses. Three recent clinical studies show reliable applications in stroke patients. After observing good test-retest reliability [110], Rogers et al. used the NeuroSky MindWave to determine distinct EEG profiles for patients who have experienced transient ischemic attack, ischemic stroke, or neither [111]; Wilkinson et al. [112] used the interaXon Muse to accurately identify patients with large vessel occlusions, a diagnostic predictor of acute stroke; and Ishaque et al. [113] used the interaXon Muse to characterise neuronal function and clinical recovery of stroke patients post-treatment.

Recording abnormal neural responses to stimuli can further assist in creating clinical profiles. Terracciano et al. [114] designed smart glasses configured with an OpenBCI Cyton board to generate a distinct visual checkerboard pattern that elicits the pattern-reversal visual-evoked potential in neurotypical populations. The onset of this component is significantly delayed in multiple sclerosis and traumatic brain injury [115]. Consumer-grade devices coupled with diagnostic tools have the potential to make accurate diagnoses more accessible for clinicians, evident from studies on insomnia [116], attention-deficit hyperactivity disorder [117], and epilepsy [118].

Seizure detection is a particularly promising application for consumer-grade EEG [119–121]. Only a single electrode is needed for detection after initial non-EEG neuroimaging localizes the most frequent source of a patient’s seizures [122]. This is achieved by using algorithms that are trained on a patient’s profile. With a portable headband or headset, epileptic patients can use affordable early detection systems, warning them and others to move to a safe place prior to seizure onset.

Lastly, a potential clinical application for consumer-grade EEG is in neurofeedback therapy. Neurofeedback is a form of operant conditioning that involves showing a patient their neural signals in real time to reinforce optimal and extinguish negative mental states [123]. Using consumer-grade devices, various neurofeedback systems have been designed for the treatment of Parkinson’s disease [13], central neuropathic pain [124,125], attention-deficit hyperactivity disorder [126,127], dyslexia [128], and depression [129].

### Validation

Validation studies comprised 6.18% of included articles in this review. The most informative of these involved comparing recordings between consumer and research-grade devices. Grummett et al. [29] found that the Emotiv had equal sensitivity to a research-grade g.HIamp in measuring SSVEPs and eyes-closed, baseline responses; although the Emotiv displayed significantly noisier spectra than the g.HIamp. Kotowski et al. [130] reported reliable measurement of the early-posterior negativity when participants responded to different emotional stimuli. Multiple studies comparing the Emotiv to a Neuroscan system provide evidence that the Emotiv performs remarkably well at recording research-grade ERPs in multi-trial experiments. De Lissa et al. [131] observed reliable recordings of the face-sensitive N170 component. Badcock et al. [132] found that the Emotiv can reliably record a series of late auditory ERP components, P100, N100, P200, N200 and P300. Barham et al. [133] found that N200 and P300 did not differ between the two systems - though, they also found that significantly more trials were rejected from the Emotiv during preprocessing. Finally, Williams et al. [134] found reliable P300, SSVEP, and mismatch negativity components. Additionally, Ries et al. [135] found that the Emotiv was comparable to a research-grade BioSemi ActiveTwo after correcting for low frequencies. In terms of event timing, Hairston et al. [136] observed only slight jitter and delay in the Emotiv’s serially-ported event timing, while Williams et al. [137] recorded comparable event timing between Emotiv and a Neuroscan.

Only a single study reported disparities between the waveforms of the Emotiv and a medical-grade Advanced Neuro Technology system: Duvinage et al. [138] found differences in their recording of P300 responses. However, as might be expected, consumer-grade devices display considerably poorer signal-to-noise ratios than their research-grade counterparts. Mahdid et al. [139] found that the functional connectivity of both Emotiv and OpenBCI systems compared poorly to research-grade systems; Raduntz [140] found that the signal reliability of Emotiv was poor if the device did not exactly fit the participant’s head; and Ekandem et al. [141] observed that the signal quality of an Emotiv (specifically, the EPOC+) declines over time as it uses a built-in rechargeable battery.

There were few validation studies for the remaining devices. Two studies compared the NeuroSky MindWave’s performance to research-grade systems. Johnstone et al. [142] found very minor differences in signal quality between a NeuroSky MindWave and a Nuamps Neuroscan; and Rogers et al. [110] found good test-retest reliability in eyes-closed paradigms, with slightly lower reliability in eyes-open paradigms. Frey [143] compared the OpenBCI’s spectral and temporal features to a medical-grade g.USBamp and found negligible differences. No validation studies were found for the MyndPlay Mindband or the interaXon Muse.

### Research location

Figure 2 shows a geographical heatmap of the number of publications by each country. In sum, the 916 studies that used consumer-grade EEG since 2010 were conducted across 83 countries spread across six continents. The top ten countries accounted for 46.72% of included studies. In order, these are: India (9.72%), United States (8.73%), China (4.48%), Malaysia (4.04%), Poland (3.82%), Indonesia (3.49%), United Kingdom (3.49%), Pakistan (3.06%), Mexico (2.95%), and Turkey (2.95%).

## Discussion

The aim of this scoping review was to explore the extent to which consumer-grade EEG devices have been applied to research domains in locations around the world. We identified 916 peer-reviewed studies that used consumer-grade EEG to collect neural data from human participants. We expand on device-specific information, location, and usage across research domains in turn.

Regarding **consumer-grade EEG devices**, Emotiv devices were the most widely used. We speculate that this was primarily due to its easy setup, relative comfort compared to other devices, and capacity to record quality ERPs. Emotiv devices require no engineering or technical expertise and configuring event markers is relatively easy using a serial port cable. However, there are a few caveats researchers should be aware of when using Emotiv. Firstly, the device has quite a low noise floor. We do not recommend using Emotiv for single-trial experiments. We also advise against using Emotiv in studies that require localising activity (e.g., functional connectivity experiments), as the plastic arms do not accommodate every head shape or size. Beyond these caveats, the Emotiv has proven useful in experiments that involve multiple trials – i.e., psychophysical or cognitive tasks. By applying task-appropriate algorithms, users can compensate for the Emotiv’s low noise floor, and the device can be implemented in BCI systems.

Despite only having a single electrode, the NeuroSky MindWave has also been widely used for research. The NeuroSky MindWave was particularly popular in BCI systems, market research, and gamification. However, there is conflicting evidence for its utility. Maskeliunas et al. [144] found that the MindWave displays poor accuracy at recognising the meditative and attentive states it claims to have built-in metrics for. Meanwhile, Rogers et al. [145] found that the device can assist in predicting functional outcomes after stroke. Thus, the NeuroSky MindWave may be useful in experiments that strictly examine the time-course of recordings from FP1 as its signal quality has been shown to be equal to or even greater than an Emotiv device. Further, the MindWave is an affordable EEG device for clinicians to complement data from other diagnostic metrics. These are rich avenues for future use of the MindWave as a research tool.

As for the three remaining devices, it is evident that more research is required to explore and validate their further experimental use. While OpenBCI has contributed to studies in engineering and robotics, more experiments are needed for an informed evaluation of OpenBCI’s applicability to cognitive neuroscience. Both the interaXon Muse and the MyndPlay Mindband have already been used in general experimental and clinical studies. Future research may address this notable gap in the literature by comparing these under-utilised devices to research-grade devices. Such studies can present the advantages and challenges associated with each device and pave the way for better-informed, specialised application of consumer-grade EEG.

Focusing now on **research domains**, this review revealed that the primary research-related use for consumer-grade EEG is in BCI systems. This may be due to the relatively low cost of administering large-*N* studies with the devices. Additionally, given enough training data, probabilistic models can now be configured for an individual output signal with impressive accuracy [146,147]. With more powerful computers and better classification algorithms, we expect that this application will only continue to grow as neuroscientists increasingly use machine learning to solve problems.

The second most research-related use was for general experimental research, spanning from cognitive tasks and recording athletic performance to low-level auditory or visual perception studies. This is further evidence that the scope of consumer-grade EEG includes a wide range of fields. Perhaps future research may focus on adding to the relatively low number of validation and signal processing studies, as these are crucial in informing users of the devices’ performance across different contexts. We also highlight the potential for consumer-grade devices to be applied to clinical studies. We observe that BCI research has provided ample evidence of the devices’ proficiency in creating biometric profiles for users, as well as prototypes for artificial robotic limbs, drowsiness detection systems, and mobility aides. With the advent of further clinical research, we may see more frequent use of the devices in hospitals and clinics, where neural augmentation has the potential to improve patient identification, diagnosis, and treatment outcomes.

As well as providing information about the scope of consumer-grade EEG devices, research domains, and research location, we note that the studies included in this review revealed an interesting trend in terms of date of publication: The number of publications that used consumer-grade EEG devices peaked in 2018, then dropped slightly in 2019 the year before the COVID-19 pandemic. During the peak years of the pandemic in 2020 and 2021, this number dropped substantially (see Figs 2C and D). This is unsurprising, as researchers’ ability to administer EEG studies to human participants in a laboratory was adversely affected by the imposition of lockdowns and travel restrictions worldwide [148–150]. While the number of studies using consumer-grade EEG was reduced during the pandemic, few EEG studies continued globally in diagnostic COVID-19 experiments [151–153]. If consumer-grade EEG devices were more well-recognised by the wider scientific community, the devices may have found utility in large-scale EEG experiments during a time when such neural data could augment epidemiological studies [154]. They may also have proved useful to cutting costs associated with the usage and maintenance of research-grade machines.

Regarding **research location**, it was encouraging to discover that the use of consumer-grade EEG is not limited to Western, industrialised countries [155,156]. We found that studies were generally reasonably spread across multiple countries, which further highlights consumer-grade devices’ portability and affordability. The same consumer-grade device used across different locations can record with roughly equivalent quality, which increases the reliability of replication EEG studies.

A core limitation of our review is the use of five databases: unlisted articles would have been missed. As such, we may not have included the full scope of studies that used consumer-grade EEG. We also only included articles published in English, and articles referring to their respective consumer-grade EEG device by a different name would also have been missed.

## Conclusion

Despite being marketed for commercial purposes, we present evidence that consumer-grade EEG has found considerable utility in scientific research. These devices have been used in non-traditional research settings and applied in innovative ways. In this paper, we provide a structured review of the current state of the literature and provide some general guidelines for their scope of use, aggregated from a large subset of consumer-grade EEG studies. From these observations, we encourage future research into more affordable and accessible neuroscientific solutions. This is a particularly salient issue given our increasingly technologically augmented lifestyles. Indeed, it is not outside the realm of possibility to see EEG being used in everyday life within the next few decades. Thus, to help inform scientists, practitioners, and the general public on the appropriate use of consumer-grade EEG, we prompt researchers to further explore the capabilities of this technology.

## Acknowledgments

JS, NW, GM, and NB designed the study. JS, NW, GM, and NB wrote the manuscript. We thank GM, NW, and NB for their useful comments and editing. JS collected the data and screened the publications. The full database of 916 included articles can be accessed with request from the authors.

This project was started with a partnership grant between Emotiv Inc. and Macquarie University. Author NW has since been employed by Emotiv. The other authors have no affiliation with Emotiv. None of the five manufacturers – Emotiv, NeuroSky, OpenBCI, interaXon, MyndPlay – were involved in the development of this manuscript.

